# Estimation of cochlear frequency selectivity using a convolution model of forward-masked compound action potentials

**DOI:** 10.1101/2022.04.15.487700

**Authors:** François Deloche, Satyabrata Parida, Andrew Sivaprakasam, Michael G. Heinz

## Abstract

Frequency selectivity is a fundamental property of the peripheral auditory system; however, the invasiveness of auditory nerve (AN) experiments limits its study in the human ear. Compound action potentials (CAPs) associated with forward-masking have been suggested as an alternative means to assess cochlear frequency selectivity. Previous methods relied on an empirical comparison of AN and CAP tuning curves in animal models, arguably not taking full advantage of the information contained in forward-masked CAPs. In this work, we seek to provide a direct estimate of the quality factor characterizing AN frequency tuning using many forward-masked CAP responses. The method is based on a convolution model of the CAP that takes into account the masking of AN populations induced by notched-noise maskers with various notch widths and attenuations. The model produces masking patterns that, once convolved by a unitary response, predict forward-masked CAP waveforms. Model parameters, including those characterizing frequency selectivity, are fine-tuned by minimizing waveform prediction errors across the different masking conditions, yielding robust estimates. The method was applied to click-evoked CAPs at the round window of anesthetized chinchillas. The estimated quality factor Q10 as a function of center frequency is shown to closely match the average quality factor obtained from AN-fiber tuning curves, without the need for an empirical correction factor. Beyond the estimation of frequency selectivity, the proposed model proves to be accurate in predicting forward-masked CAP responses, and therefore could be extended to study more complex aspects of cochlear signal processing using a similar experimental approach.

## Introduction

Much of our knowledge about the mammalian peripheral auditory system has been gained from single-fiber recordings of the auditory nerve. However, the invasiveness of these experiments prevents their use in humans, hampering the search of potential specificities of the human auditory system. Other means have been employed to infer the properties of the human inner ear, either through psychophysical experiments, or through less invasive physiological methods. In particular, a combination of these solutions – including psychophysical experiments based on masking [Shera et al., 2002, Oxenham and Shera, 2003], otoacoustic emissions (OAEs) [Shera et al., 2002, Sumner et al., 2018] and compound action potentials (CAPs) [Verschooten et al., 2018] – has led to a growing body of evidence that cochlear frequency selectivity is sharper in humans than small mammals. Frequency selectivity is a fundamental property of the peripheral auditory system, but its study is not straightforward, a reason being that it is affected by cochlear compressive nonlinearities [Heinz et al., 2002, Oxenham and Shera, 2003, Eustaquio-Martín and Lopez-Poveda, 2011]. As a result, although data on cochlear frequency tuning in humans have been obtained by various means, the picture is not as detailed as for other mammals, and some methods of assessing cochlear frequency selectivity do not show any significant difference with small mammals [Ruggero and Temchin, 2005]. To advance our knowledge in this area, it is necessary to refine the available tools and to better understand how they relate to auditory physiology. For example, OAE-based estimates of frequency selectivity would benefit from a better understanding of how OAE delays [Shera and Charaziak, 2019] or distorsion-product level functions [Wilson et al., 2021] relate to cochlear tuning. The focus of this paper is the compound action potential (CAP), an auditory evoked potential that reflects the summed activity of auditory nerve fibers (ANFs). CAP data can be obtained with a satisfactory signal-to-noise ratio (SNR) at the cost of moderate invasiveness [Eggermont, 2017, Verschooten and Joris, 2022], and, if analyzed with an appropriate model, could provide a lot of information on the compound response of ANFs.

Methods of frequency selectivity estimation based on the CAP rely on the masking paradigm, similar to psychophysical experiments historically associated with the measurement of critical bands in humans [Patterson, 1976, Moore and Glasberg, 1983]. While simultaneous masking reflects both excitatory and suppressive masking [Delgutte, 1990, Harrison et al., 1981b, Charaziak and Siegel, 2014], estimates based on forward masking reflect only excitatory masking and have good agreement with ANF tuning curves [Harrison et al., 1981a,b, Verschooten et al., 2012]. In the last decade, Verschooten et al. refined a previous estimation procedure based on forward-masked CAPs [Harrison et al., 1981a,b] using the notched-noise method. The advantage of notched-noise maskers over narrowband stimuli is that results are less confounded by suppression effects; they also limit the effect of ‘off-frequency listening’ [Delgutte, 1990, Oxenham and Shera, 2003], i.e., the activity of ANFs tuned at frequencies below or above the notch is reduced. The procedure of Verschooten et al. was first validated in animal models [Verschooten et al., 2012] and later applied to human subjects [Verschooten et al., 2018]. Their estimation method was based on establishing iso-response curves for masker level versus masker notch width. However, the method required an empirical correction factor to match the quality factor *Q*_10_ of ANF tuning curves, which was not the same for every species. In particular, a higher estimate of human cochlear frequency tuning was obtained when the correction factor found for macaques was applied, leaving the exact range for *Q*_10_ uncertain.

In this work, we propose a method that seeks to estimate the frequency tuning of ANFs directly, avoiding the need for an empirical correction factor. To this end, we assume that the masked part of forward-masked CAP responses can be approximated by a ‘masking pattern’ convolved by a unitary response. Convolution models for the CAP have been used for decades [Goldstein and Kiang, 1958] but with limited applications, since this type of model requires many assumptions about the multiple factors that affect CAP waveforms, in particular: the (level-dependent) relationship between cochlear place and AN spike latencies, the spread of excitation along the cochlear partition, the unitary response and the distribution of thresholds and rate functions [Boer, 1975]. Considering the forward-masking of a CAP response in multiple different masking settings but with a fixed probe appears to be a less challenging option from the modeling perspective, given in addition that masking reveals information about the different factors mentioned (e.g., the place-latency relationship using high-pass noise maskers [Eggermont, 1976]). In this paper, we introduce a differentiable model for predicting the waveforms of forward-masked click-evoked CAPs when presented with notched-noise maskers. The model is applied to CAPs recorded at the round window of anesthetized chinchillas, and the estimation of model parameters based on the minimization of the prediction error by gradient descent is assessed. In particular, we show that the resulting estimates for the quality factor provide an excellent match to published ANF tuning curve values.

## Methods

### Experimental setting

#### Preparation & Anesthesia

CAP responses were acquired in 5 adult male chinchillas (*Chinchilla lanigera*) using surgical procedures pre-approved by the Purdue Animal Care and Use Committee. Anesthesia was induced using subcutaneous injections of xylazine (2-3 mg/kg) and ketamine (30-40 mg/kg). Anesthesia was maintained using intraperitoneal boluses of sodium pentobarbital (15 mg/kg/2h), and fluids (Lactated Ringer’s) were administered subcutaneously throughout the experiment (~1cc/hr). The animals’ vital signs were monitored throughout experiments using pulse oximetry (Nonin 8600V, Plymouth, MN) while oxygen was continuously delivered to the animal. Body temperature was maintained at 37°C using a homeothermic monitoring system with rectal probe (50-7220F, Harvard Apparatus).

#### Surgical Procedure

Following anesthetic induction, a tracheotomy was performed to provide a low-resistance airway, reducing respiratory artifacts. Skin and muscles were transected following a dorsal-midline incision, and the external ear canals and bullae were subsequently exposed. Hollow ear bars were bilaterally placed in the ear canals and secured to a stereotaxic frame (David Kopf Instruments, Tujunga, CA). Sound was delivered monaurally through the ear bars using a dynamic loudspeaker (DT48, Beyerdynamic). To prevent a progressive negative pressure buildup in the bulla, a polyethylene tube (PE-90) was placed through an incision in the anterior bulla [Guinan and Peake, 1967]. A second incision was made in the posterior base of the ipsilateral bulla to expose the middle ear. A silver wire electrode was placed near the round window to record CAPs and sealed in place within bulla opening using light-cured dental cement (Prime-Dent, USA). A pocket in the nape of the neck was made for a silver coiled wire reference electrode soaked in isotonic saline and connected to ground. All procedures were carried out in a double-walled, electrically shielded, sound-attenuating booth (Acoustic Systems, Austin, TX, USA). At the end of the experiments, animals were euthanized by barbiturate overdose.

#### Signal acquisition and pre-processing

We calibrated sound input using a probe microphone (Etymotic ER-7C) placed near eardrum. A flat frequency response (within ± 2 dB until 10kHz) was achieved using a 256-tap digital finite impulse response filter for the forward-maskers. For the click probe, we adopted a different strategy by using the inverse of a 128th-order all-pole filter computed using linear predictive coding (LPC) to correct for the phase differences induced by the acoustic system. CAP responses from the round window were amplified and band-passed using an ISO-80 Bio-Amplifier (10^3^ gain, 10^2^ — 10^4^ Hz, World Precision Instruments) before being recorded by hardware modules from Tucker-Davis Technologies (TDT, Alachua, FL). Signal acquisition was controlled by a custom MATLAB-based (MathWorks, Natick, MA) interface. Of the 5 chinchillas tested, 4 had exploitable data at all center frequencies (CFs) tested (except at CF=8 kHz for chinchilla Q333). Masking had a too small effect on the CAP response for the remaining animal (signal-to-noise ratio too low or absence of signal for Δ*CAP*), except for CFs in the range 3–5 kHz. This animal is not included in the Results section, but the analysis we conducted on the partial data did not contradict the results obtained for the other animals. Prior to analysis, the CAP responses were pre-processed by applying a Tukey window to keep only the part where most of the masking occurred (typical parameters: window length 4 ms, proportion of interval covered by the tapered cosine region: 0.4). The signals were smoothed by a Gaussian filter of standard deviation 0.03 ms. Specific experimental artifacts were addressed by additional pre-processing in two animals: correction of a DC drift and removal of a periodic noise using a notch filter.

### Stimulus paradigm

#### Presentation of masker and probe

CAPs in response to alternating-polarity 80 dB peSPL clicks were recorded from the round window in the presence of Gaussian noise maskers according to a forward-masking paradigm. Fig 1 A shows the time representation of the masker and probe, and the durations within one stimulus cycle, with one cycle totaling 160 ms. We used in total around 150 masking conditions, each associated with a specific power spectrum profile (Fig 1 B). The masking conditions are further described in the next paragraph. Each condition was repeated 120×2 times (12 blocks × 10 repetitions × 2 click polarities). Within each block, the conditions were presented in a random order, ensuring some degree of interleaving. The interleaving of conditions was important to avoid unintended effects of long-term adaptation. Such effects on the CAP responses were indeed visible in a pilot experiment without interleaving constraint, especially 2.5 ms after the onset of the CAP (N2), possibly corresponding to the response from the cochlear nucleus [Møller, 1983].

**Figure 1:**
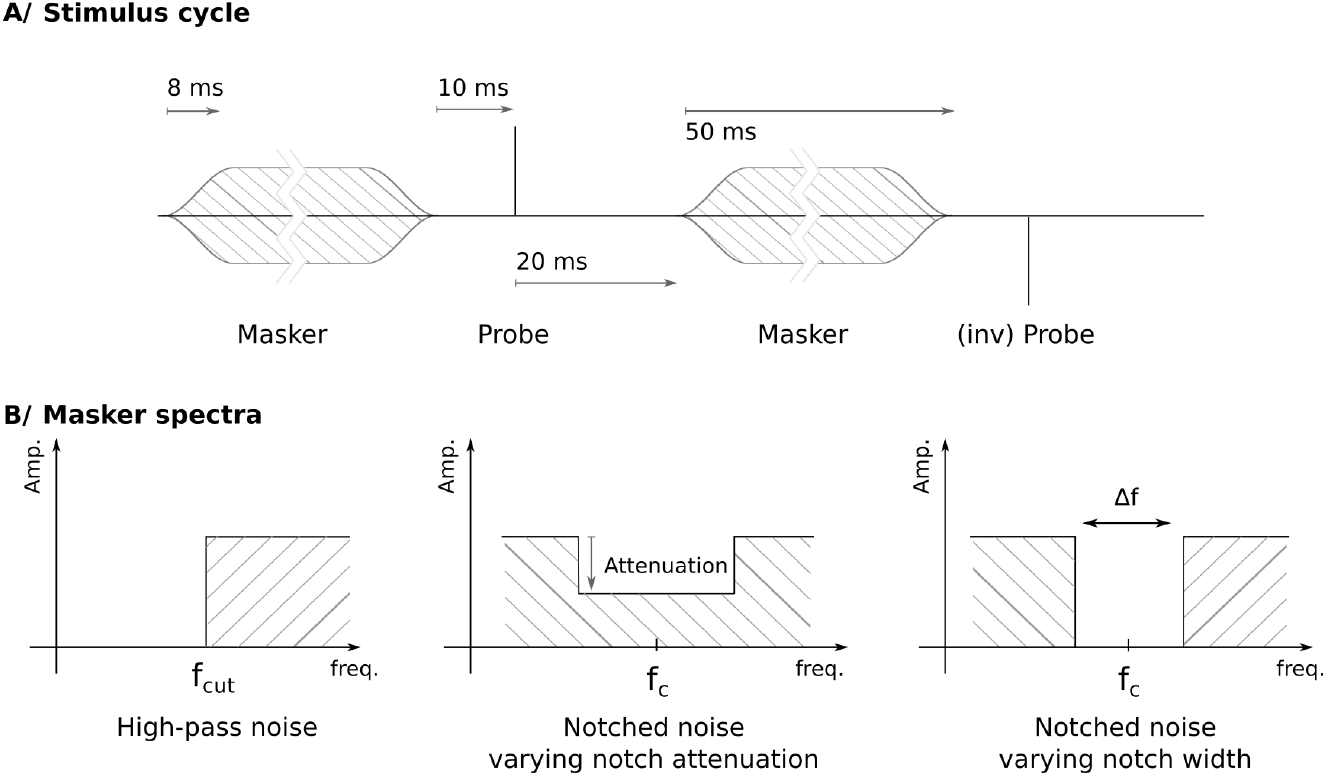
Time representation of one stimulus cycle (**A/**) and spectral representation of the three types of maskers (**B/**). **A/** The stimuli consisted of the repetition of a masker and probe. The masker was generated from Gaussian noise with a spectrum that follows a pattern represented in panel B/. The probe was a click of alternating polarity. The forward-masked CAPs were obtained by averaging the responses evoked by the probe presented under the same masking condition. The durations illustrated are from left to right: gating time (cosine ramp), masker-probe interval, probe-masker interval and masker duration. **B/** Schematic representation of the spectra of the three different types of maskers. Each type of masker was designed for a different purpose: high-pass noise maskers for the estimation of the place-latency relationship (‘narrow-band analysis’ method [Eggermont, 1976]), notched noise maskers for the estimation of masking input-output functions (maskers with varying notch attenuation) and frequency tuning (varying notch widths).

#### Masker design

The 155 maskers were of three different types, corresponding to high-pass noise (12 maskers), notched noise with varying amplitude for the notch (77 maskers), or notched noise with a varying notch width (65 maskers). This set also includes the broadband noise masking condition (no notch), used as the reference condition. The high-pass noise maskers were used to reproduce the narrow-band analysis of the CAP [Eggermont, 1976, Prijs and Eggermont, 1981] that provides estimates of the latencies associated with the cut-off frequencies of the maskers. The latencies follow the same trend as the cochlear traveling wave, with the onset of basal contributions (high CFs) preceding the onset of apical contributions (lower CFs) – e.g. in our data, contributions corresponding to 2 kHz are delayed by 0.5 ms compared to the most basal frequencies. The 12 cut-off frequencies associated with the high-pass maskers ranged from 1.2 kHz to 10 kHz. The two other types of masking conditions were notched-noise maskers with the notch having either a varying amplitude or a varying notch width. The notched-noise maskers were grouped according to the center frequency of the notch which was around 7 reference frequencies: 1.5, 2.2, 3, 4, 5, 6, and 8 kHz. For each CF, 10 maskers corresponded to the varying notch amplitude type, ranging from 35 dB attenuation to 0 dB attenuation, thus gradually merging into the broadband noise condition (0 dB attenuation, reference condition). Except for the first experiment that was conducted (chinchilla Q395), an additional condition was included corresponding to a notched-noise masker with −3 dB attenuation for the notch (i.e., the power spectrum in the region of the notch was above the broadband noise spectrum density); the introduction of this extra masker helped to determine the slope of input-output masking curves at the reference point. The notched-noise maskers with different notch widths typically had a large notch width (e.g., 2 kHz at CF=5 kHz, 1 kHz at CF=1.5 kHz). They were used to estimate the amount of masking as a function of place-specific response intensity (input/output masking curves, see further in text). Around 10 maskers by CF were related to the last type of maskers: notched-noise maskers with a varying notch width. As an example, the 10 maskers associated with this type at CF=5 kHz had a notch width in the range 900 Hz to 1.4 kHz, which is of the order of the expected value of the 10-dB bandwidth of cochlear filters at this CF [Temchin et al., 2008]. The center frequency of the notch was allowed to vary slightly to probe different groups of ANFs, e.g. 4 800 Hz for one masker and 5 200 Hz for another. The notch amplitude for this type of masker was in most cases zero. These maskers were designed to estimate the frequency selectivity of the auditory filters. The rationale is the same as for psychological measurements of critical bands using notched noise stimuli [Patterson, 1976, Moore and Glasberg, 1983, Oxenham and Shera, 2003], and derives from the following principle: if increasing the notch width from the broadband condition results in a significant masking release effect, it indicates a sharp tuning of cochlear filters. The frequency spectra of all the maskers were restricted to the range between 200 Hz and 12 kHz and the power spectral density (PSD), excluding the notches and filtered parts, was set to a constant. For all the animals, the maximum PSD was similar, in the range 4–14 dB SPL, corresponding to a sound level in the range 45–55 dB SPL for the broadband noise condition.

### Model

#### Main hypothesis

The main hypothesis of our model is that the masking of the CAP is driven by the response intensity at the output of a cochlear filter bank and input/output (I/O) masking curves determining the growth of masking. More explicitly, if *I* is the average intensity in response to the masker at the output of a cochlear filter, we assume that the amount of masking *M* for the compound response of the associated ANFs can be represented by a function of *I*. We tested two common functions for these I/O masking curves (shown in Fig 4 A in the Results section):

- the sigmoid:

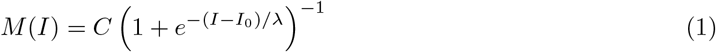
- the Weibull cumulative distribution function (CDF), as in [Verschooten et al., 2012]:

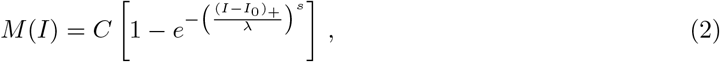

where (*I* — *I*_0_)_+_ = *I* – *I*_0_ if *I* ≥ *I*_0_, 0 elsewhere.

The unknown parameters for the sigmoid are *I*_0_ and the scale parameter *λ*. The additional shape parameter *s* allows the Weibull CDF to fit a larger set of functions that do not necessarily have symmetry around their half-maximum value point. By convention, we set the constant *C* so that the I/O masking functions are constrained to 100% masking for the response level corresponding to the broadband noise condition (reference level).

#### Generation of the CAP estimates

The model builds on the convolution model that was already used in early work on the CAP [Goldstein and Kiang, 1958, Boer, 1975]. Conveniently, the convolution model for the CAP can be written as a convolution between a cochlear excitation pattern *E* defined in the latency domain and a unitary response *u*_0_:

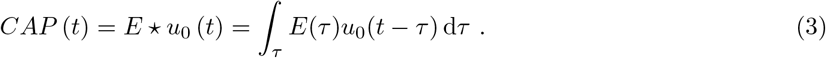

In this form, *u*_0_ accounts for the spike unit response but also for the spike histogram of a population of synchronized ANFs normalized with respect to the number of spikes (see [Elberling and Hoke, 1978] or [Bappert et al., 1980] for a similar approach). The decomposition of *u*_0_ under the hypothesis of a constant normalized spike histogram – justified in particular if we consider a narrow range of cochlear locations – in Eq 3 leads to a *double* convolution model:

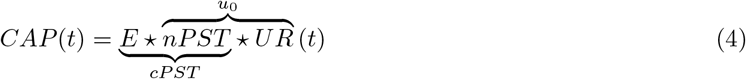

where *nPST, cPST* stand for the normalized and compound post-stimulus time histograms (PSTH), and *UR* is the spike unit response.

The focus of our method, instead, is the masking of the CAP. *CAP*(*t*) is therefore replaced by Δ*CAP*(*t*), the release of masking of the CAP, defined by the difference in amplitude between the CAP response when a notched-noise masker is presented versus the response corresponding to the broadband noise masker:

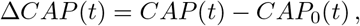

where *CAP*_0_ is the reference response with the broadband noise condition as masker.

An example of how Δ*CAP*(*t*) is derived from *CAP*(*t*) with typical data is shown in Fig 3 in the Results section. We can write a similar equation to Eq 4 for Δ*CAP*:

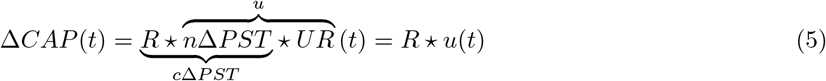

where *R*(*τ*) is the *masking release pattern* and *u* is the unitary response. Again, *u* is the compound of the spike unit response and the difference in the PSTH of a population of synchronized ANFs normalized with respect to the amount of masking (*n*Δ*PST*). We are not interested, however, in the exact decomposition of *u* and we will refer to the simpler equation Δ*CAP* = *R*u* in the rest of the paper. Note that the model assumes that *n*Δ*PST* is invariant regardless of the amount of masking. Prior to any experiment, we tested whether this hypothesis was reasonable with a well-established computational model of ANF responses [Bruce et al., 2018]. This analysis is left as supplementary material (*SI_PSTH.pdf*). As for the spike unitary response, authors reported that it can be essentially considered independent of the ANF best frequency or spontaneous rate [Kiang et al., 1976, Wang, 1979, Prijs, 1986].

Figure 2 shows the steps leading to the generation of the estimates 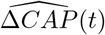. In the following, we describe and justify these steps going backward from 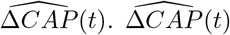 is obtained by convolution of a masking release pattern and the unitary response (Eq 5). We consider that the masking release pattern *R* is related to the amount of masking *M* by

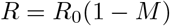

where *R*_0_ represents the relative contributions of ANF populations to Δ*CAP*. *R*_0_ can also be seen as the masking release pattern when there is no forward-masker. By convention, *M* = 1 for the broadband noise condition. *R*_0_, *M* and *R* depend on latency (e.g. *R* = *R*(*τ*)), but we consider that place (i.e., center frequency) and latency are related by a power-law: 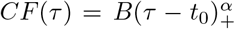, where *B* and *α* are estimated using the high-pass noise maskers (following the narrow-band analysis method [Eggermont, 2017]). Hence, all latency dependencies can be converted to a place dependency and vice versa. For the generation of the masking release patterns, we estimated *R*_0_ and *M* first in the frequency domain then the masking release pattern was converted to the time domain. *R_0_*(*f*) can be considered as frequency weights, which have to be included to account for the non-homogeneous contributions of different CFs to the masking release of the CAP. Finally, to compute the amount of masking *M* as a function of frequency, we relied on a simplified model of cochlear filtering using a linear filter bank. Given the average power spectral density of the masker *S*(*f*), the average response intensity at the output of a cochlear filter characterized by CF was computed using:

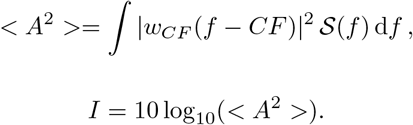

**Figure 2:**
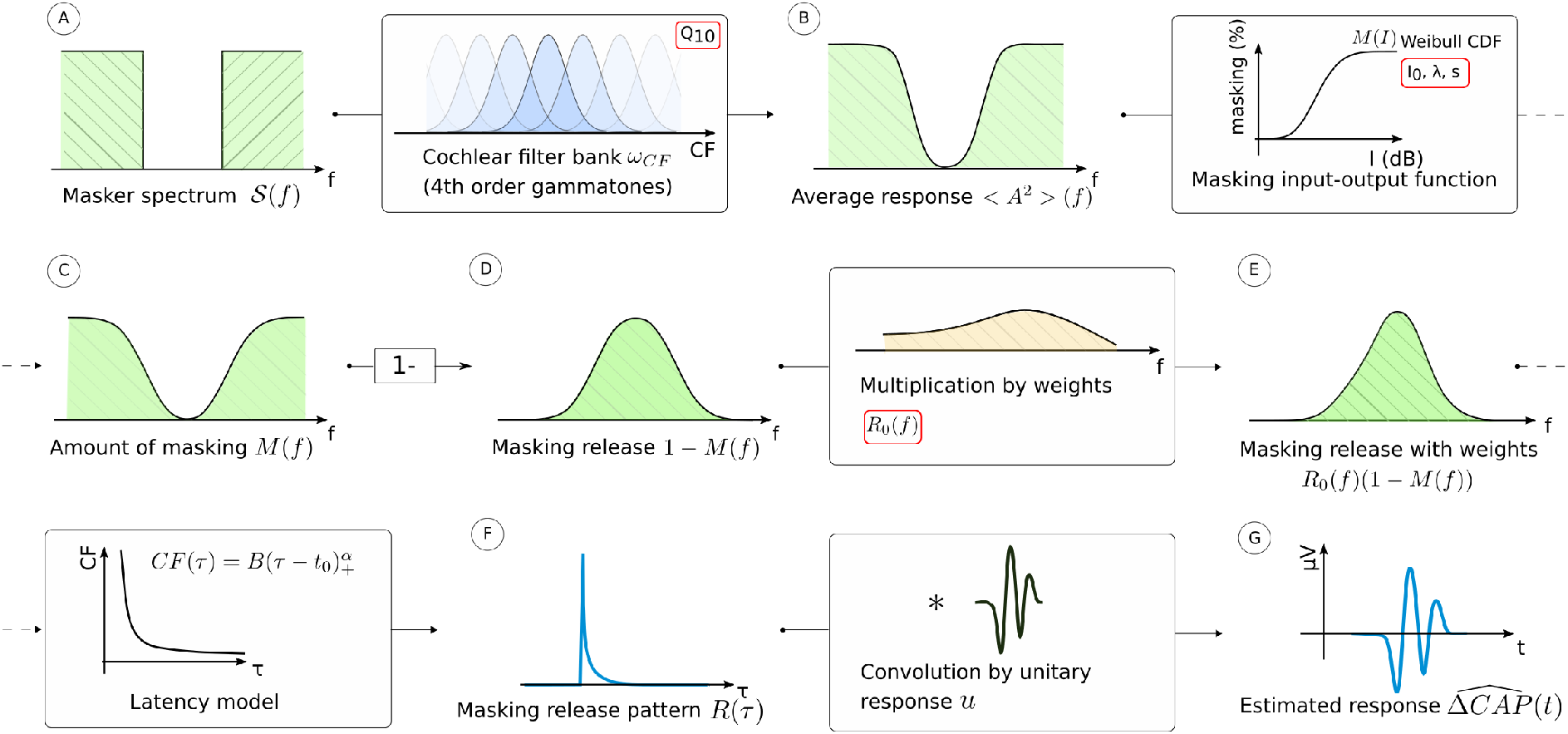
Flow diagram of the generation of the masking release estimates 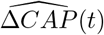. The masker spectrum (**A**) is provided at the input and decomposed by a model of cochlear filter bank with gammatone filters. As the masker spectra are of simple form, i.e., composed of rectangular bands, the average response (**B**) at the output of the filter bank was computed using an analytical formula (see text). The masking input-output function applied to the average response provides the amount of masking *M*(*f*) (**C**) or, equivalently, the amount of masking release 1 — *M*(*f*) (**D**). Frequency weights *R*_0_ (*f*) are included to the result to account for the non-homogeneous contributions of different frequencies to Δ*CAP*. This yields the final estimate of the amount of masking release defined in the frequency domain (**E**). Using a power-law model for the latencies, the masking release is converted to the time domain, giving the masking release pattern *R*(*τ*) (**F**). Once convolved with the unitary response *u*, we finally obtain the estimate of the release 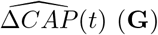 (**G**). The parameters that are fine-tuned during the optimization process (gradient descent) are highlighted in red: they are *Q*_10_, the masking I/O function (Weibull CDF) variables, and the frequency weights. The unitary response *u* and the power-law parameters relating CFs and latencies are also parameters of the model, but are adjusted independently before the optimization procedure.

The amount of masking *M* was then obtained by applying the I/O masking function (Eq 1 or 2) on *I*. *w_CF_* is the cochlear filter shape in the frequency domain, defined such as its root mean square (RMS) value is 1. We implemented two models of cochlear filters for *w_CF_*: gammatones (as illustrated in Fig 2 and used for the Results section) and Gaussian filters. Once the type of cochlear filter is chosen, *w_CF_* depends only on the tuning of the cochlear filter at CF, characterized by the quality factor *Q*_10_ (related to the 10 dB-bandwidth by: *Q*_10_ = *CF/BW*_10_). As the masker spectra are simple and defined by rectangular bands, analytical formulas for *I* as a function of CF were employed instead of integral expressions. For instance, in the case of the Gaussian implementation and considering a masker spectrum made of a single band [*f*_low_, *f*_high_] with power *S*_0_, we have:

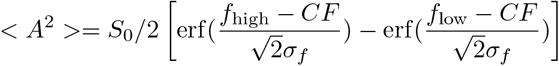

with 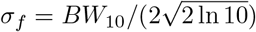 and erf being the error function. In the case of a masker presenting multiple bands, the expressions for each band simply add up. The derivation of the formula for gammatones is presented in the Appendix.

### Estimation and optimization procedure

#### Model unknowns

The model presented in the previous paragraph and outlined in Fig 2 has multiple unknowns that are reviewed here:

1. The relationship between latencies and center frequencies. It is assumed to follow a power-law: 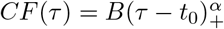.
2. The unitary response *u*.
3. The amount of masking as a function of place-specific intensity response (*I/O masking curves*). In the case of the Weibull CDF (Eq 2), as adopted in the rest of the paper, this curve is parametrically defined by three variables (*λ*, *I*_0_, *s*).
4. The tuning of the auditory filters, characterized by *Q*_10_.
5. The frequency weights *R*_0_ (*f*) characterizing the contributions of each cochlear region (or frequency interval) to Δ*CAP*.

#### Estimation procedure

While fitting all these parameters at once could seem intractable in a traditional setting, this approach is made possible by the fact that responses to many masking conditions are acquired during an experiment. It is also technically facilitated by the existence of automatic differentiation libraries (see details of the optimization procedure below). This section presents the outline of the estimation procedure. The technical details of the step-by-step procedure can be found in the code released for this project [Deloche, 2022].

The two unknowns that are determined first are the place-latency relationship and the unitary response. The relationship between latencies and center frequencies is determined using the narrow-band analysis method [Eggermont, 2017]. The method is based on the high-pass noise maskers; when presented in the order of decreasing cut-off frequencies, these maskers progressively mask the basal contributions to the CAP. The CAP peak delay is assumed to be the latency associated with the cut-off frequency. The latencies were fit by a power-law estimated from the peak delays by least-squares fitting (dog leg method). The unitary response *u* was estimated by deconvolution of the release-of-masking signals [Δ*CAP*(*t*)] with a first estimation of the masking release patterns for the notched noise maskers. For this step, the CAP responses were smoothed by a Gaussian filter of deviation 0.06 ms (instead of 0.03 ms elsewhere). Once the unitary responses and latencies were determined, they were considered fixed during the rest of the estimation procedure. However, after the optimization of the other parameters was done, the unitary response was re-estimated with the updated masking release patterns, and the optimization procedure was performed a second time.

All the other model parameters, highlighted in red in Fig 2, were fitted simultaneously by minimizing the mean squared error (MSE) between the signals 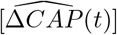 generated by the model, and the true signals [Δ*CAP*]. The optimization procedure is described in the next paragraph. One challenge of the method is that most of the model parameters potentially depend on CF. This is the case for the unitary response, the parameters controlling the I/O masking curve, the quality factor *Q*_10_, and the frequency weights *R*_0_(*f*). This issue is mostly resolved by adjusting different versions of the model to each CF probed instead of having a single model fitted on all the data. For this purpose, the notched-noise maskers were grouped into 7 different center frequencies according to the frequency region of the notch (CF = 1.5, 2.2, 3, 4, 5, 6 or 8 kHz) and their responses were fitted separately. However, we allowed some parameters to be shared across the different optimization nodes. This was in particular the case for the frequency weights *R*_0_(*f*). The estimation of *R*_0_(*f*) at every frequency was made possible at the cost of a regularity assumption. We assumed that *R*_0_ belongs to a low-dimensional manifold, explicitly that *R*_0_(*f*) in the range [200 Hz, 12kHz] is only defined by its *m* first Fourier coefficients (*m* = 10). For the estimation of *Q*_10_, we assumed that the 10-dB bandwidth was constant in the interval of frequencies around CF and searched its optimal value using gradient descent or a grid search method. As an alternative, we also used a regression method assuming that *Q*_10_ could be approximated by a radial basis function (RBF) network (6 hidden neurons, input *x* = *f*/15000, output: log *Q*_10_, activations are Gaussian functions with *σ* = 0.5). The results of the two methods are shown at the end of the paper.

#### Optimization procedure

The goal of the optimization procedure is to adjust the model parameters highlighted in red in Fig 2, including *Q*_10_ characterizing frequency selectivity, to obtain the best fit between the signals generated by the model and the true responses. We denote [Δ*CAP*(*t*)]_*i*_ the masking releases of the CAP, where *i* is an index for the masking condition (*i* = 1 …. *N*_cond_, with *N*_cond_ the number of masking conditions).

The model yields estimates 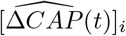 for each masking condition, and we define the cost function as the total mean square error:

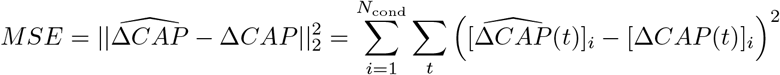

*MSE* was minimized through a gradient descent scheme. The gradients with respect to the model parameters were computed with PyTorch, an automatic differentiation library originally designed for the optimization of artificial neural networks [Paszke et al., 2019]. A schematic for the graph of computations is provided in supplementary materials *(SI_computations.pdf*), that also synthesizes the operations that lead to the generation of 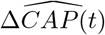. The key point is that, although the entire model is complex, each step of computation is a simple differentiable operation, and the gradients can be computed by applying the chain rule. An alternate gradient scheme was adopted. At step 1, the gradients were computed and summed over all the notched-noise masker conditions and the frequency weights *R*_0_ (*f*) were updated. At step 2, the gradients were computed over the maskers with a notch of varying amplitude and the I/O masking function was updated. At step 3, *Q*_10_ was updated using the the maskers with a varying notch width. The same steps were then repeated about 100 times. The optimization was done separately for each CF probed. However, some parameters were shared and optimized jointly (e.g. frequency weights *R*_0_(*f*)), using the distributed communication package of PyTorch. The parameters were initially set manually or set at plausible values (e.g., *Q*_10_ was set to fit the curve *Q*_10_ = 2(*f*/1000)^0.5^), before being fine-tuned by the optimization algorithm. Since the cost function is not guaranteed to be convex with respect to the model parameters, and the algorithm can be stuck in local minima, several initializations were tried.

## Results

The results presented in this section were obtained using the 4th-order gammatone model for cochlear filters and the Weibull CDF for the masking I/O functions. It can be noted, however, that we did not find significantly different results when using Gaussian filters instead of gammatones. The first figures in this section show an example of forward-masked CAP data and the ancillary model parameters in one animal (chinchilla Q395).

### Estimation of input-output masking curves

A first indication of the masking input-output curve – the amount of masking as a function of cochlear-filter output intensity – is provided by the measure of reduction of the Δ*CAP* peak amplitude when the masker presents a notch of decreasing attenuation centered at CF. Fig 4 A displays this type of data corresponding to the CAP responses shown in Fig 3. The sigmoid and the Weibull CDF equally provide a good fit for this example, but a larger number of cases were matched by the Weibull CDF, which allows for greater flexibility. In reality, the relationship between reduction of the CAP peak amplitude and the underlying masking I/O curve is not guaranteed to be linear, because the masking of the CAP also depends on the spread of the cochlear excitation pattern, which differs for each masker. For this reason, the determination of the reduction of the CAP amplitude serves only as a first approximation of the parametric I/O masking curve, which is then fine-tuned during the optimization procedure along with the other parameters. The masking I/O curve at CF=5 kHz after optimization (dashed line) is also shown in Fig 4 A, clearly deviating from the initial curve. The other curves for the same animal at different CFs are shown in panel B. We did not find a regular pattern in the changes of the I/O curves with CF considering all the animals in the study. Note that since the I/O functions were found using notched-noise maskers, the amount of 0% masking does not necessarily mean that no masking occurs for that level but rather that no masking is discernible from the masking elicited by the sides of the masker.

**Figure 3:**
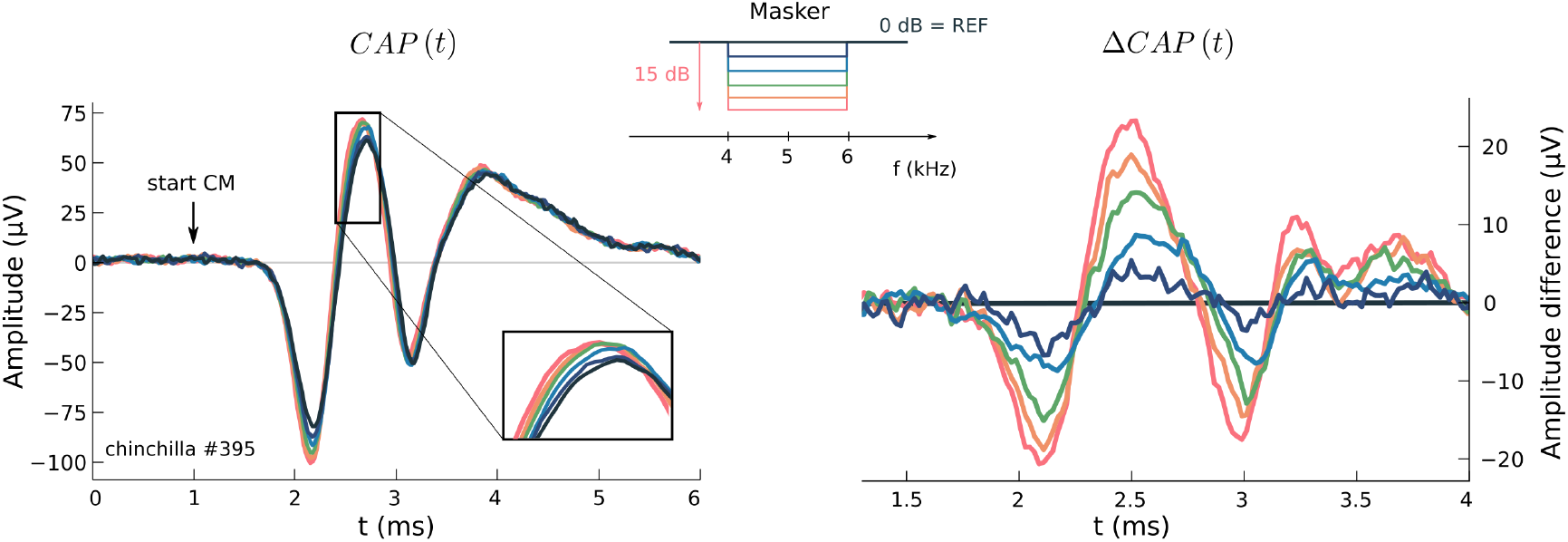
Example of CAP data and derivation of Δ*CAP*(*t*). **Left:** forward-masked CAP responses to 80-dB clicks with a Gaussian noise masker presenting a 2-kHz notch of varying amplitude around 5 kHz (masker profiles are shown at the top center). **Right:** Masking release of the same forward-masked CAPs, using the broadband noise condition as reference (Δ*CAP*(*t*) = *CAP*(*t*) — *CAP*_0_(*t*)). CM=cochlear microphonics. Notch attenuations: 15, 12, 9, 6, 3, 0dB (REF).

**Figure 4:**
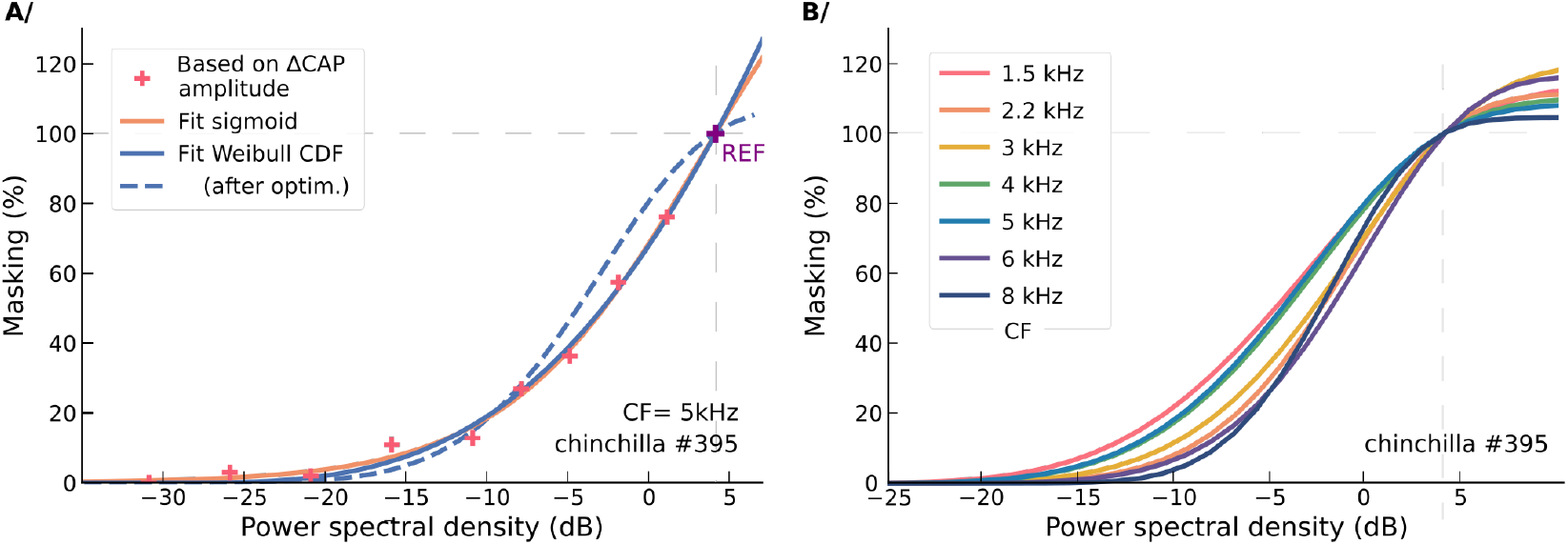
Masking input-output (I/O) curves. **A/** Amount of masking at CF=5 kHz as estimated by the peak-to-peak amplitude of the responses represented in Fig 3 (masker with a 2 kHz-wide notch centered at 5 kHz). The x-axis refers to the power spectral density within the notch. The purple cross corresponds to the reference condition (0 attenuation relative to maximum PSD, i.e., broadband noise condition, matched to 100% masking). Fits with the sigmoid and Weibull CDF functions are shown, as well as the Weibull CDF fit after fine-tuning the model (dashed line), considered to better approximate the underlying masking I/O function of the compound response of ANFs tuned to CF. **B/** Masking I/O curves (Weibull CDFs) for the same animal at all the CFs after fine-tuning the model.

### Estimation of latencies, unitary responses, frequency weights

Figure 5 shows the estimated latencies for the same chinchilla using the narrowband analysis method. Small deviations (<0.15) from the power-law can be observed. These deviations appear bigger on a log-log scale for high CF, but these do not affect the overall performance of the model since the latencies for these frequencies are in fact small. The values of the latencies are to be interpreted along with the peak delays of the estimation of the unitary response *u*, shown for the same animal in Fig 6 A. *u* keeps the biphasic shape of the spike unitary response [Wang, 1979] but is repeated at least twice, with the two first negative peaks separated by 0.8 ms. The second peak has been partly attributed to the phenomenon of ‘double-spiking’, i.e., the firing of ANFs immediately after the refractory period [Özdamar and Dallos, 1978, Versnel et al., 1992]. Another reason may be the presence of sub-threshold electrical resonances in the auditory nerve peripheral dendrites [McMahon and Patuzzi, 2002]. Interestingly, this figure does not exhibit significant variations in the shape of the unitary response, but small changes with a trend consistent with decreasing CFs can be observed at 2.2 and 3 ms. These changes could be explained by larger group delays for apical cochlear filters (i.e., a slower build-up of response intensity), hence a broader *nΔPST* for lower CFs in Equation 5. A fast analysis based on the deconvolution of the unitary responses at each CF with the unitary response at CF=8 kHz tends to confirm this hypothesis. Although we only described the results for one animal in this paragraph, the results were similar for the other animals in the study. We observed, however, larger deviations from the power-law for latencies in two animals, especially at lower CFs with deviations up to 0.3 ms at CF=1.5 kHz.

**Figure 5:**
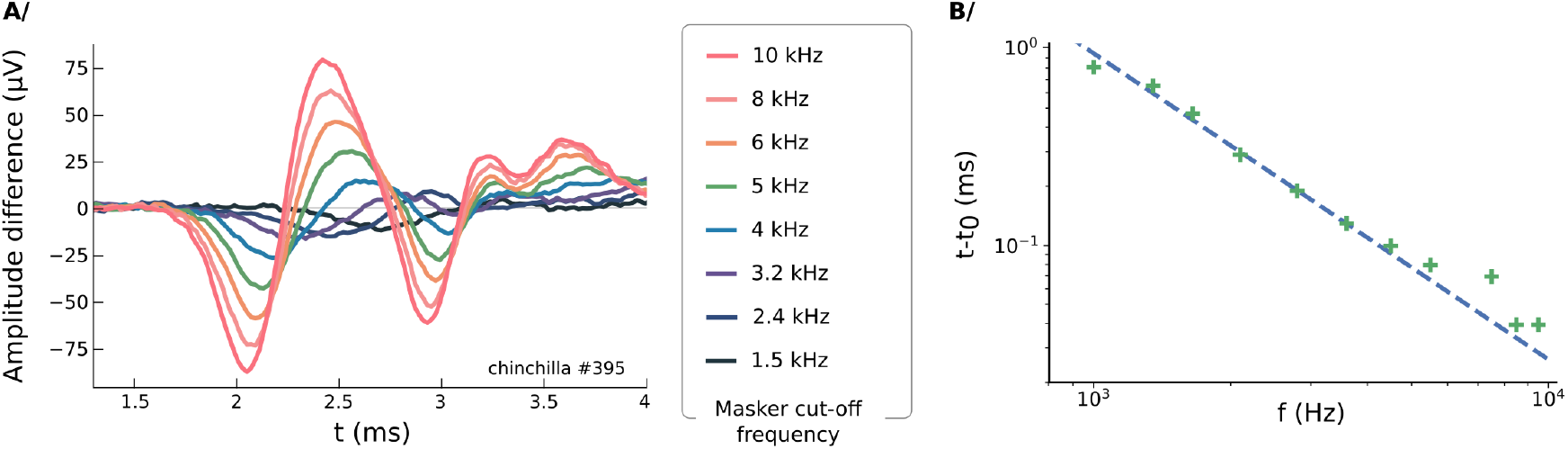
Estimation of the place-latency relationship. **A**/ Release-of-masking Δ*CAP* for the high-pass noise maskers. The cut-off frequency goes from 10 kHz to 1.5 kHz (REF: broadband noise, cut-off frequency 200 Hz). The responses display the shift of the peak latencies that reflect the cochlear traveling wave, with the CAP onset corresponding to the contributions of the most basal fibers. Responses for 8 high-pass noise maskers out of 12 are represented. **B/** Results of the estimation of the latencies as a function of frequency (green crosses, log-log scale) using the narrowband analysis method [Eggermont, 1976] with the signals represented in panel A. Fit (dashed line): power law, *f* = 11.6 (*t* – *t*_0_)^-0.64^, with *t*_0_ = 0.83 ms (to consider along with *u*, Fig 6 A), standard error: 0.05 ms.

Fig 6 B shows the distribution of weights *R*_0_ accounting for the relative contributions to Δ*CAP*, both in the place-frequency domain and in the latency domain (panel B). The estimation of the distribution *R*_0_(*f*) has been our main difficulty to fit CAP data and obtain consistent estimates. The slow decreasing trend of *R*_0_(*f*) was expected as a result of the spatial distribution of inner hair cells (i.e., the exponential relationship between cochlear place and frequency). However, as shown in Fig 6 B, *R*_0_(*f*) exhibits in addition two narrow dips (2.5 kHz and 6 kHz) that hinder the estimation, not only of the frequency weights, but also of the other parameters of the model at the corresponding frequencies. In the same time, since *R*_0_(*f*) is estimated as a sum of sine and cosine functions, oscillations in the approximation of *R*_0_(*f*) can damage the prediction of other model parameters. To deal with this issue, we adopted a strategy consisting in approximating *R*_0_(*f*) with low modes only (*m* = 4) at initialization of the optimization procedure, then increasing the maximum mode (*m* = 10) while conducting gradient descent. Most of the chinchillas presented the same type of distributions, with an overall decreasing trend for *R*_0_(*f*) and one or two relatively narrow dips, but the dips were not always found at the same frequencies. We do not have a clear explanation for the presence of dips in *R*_0_ (*f*), although one hypothesis is that they result from the three-dimensional cochlear geometry.

**Figure 6:**
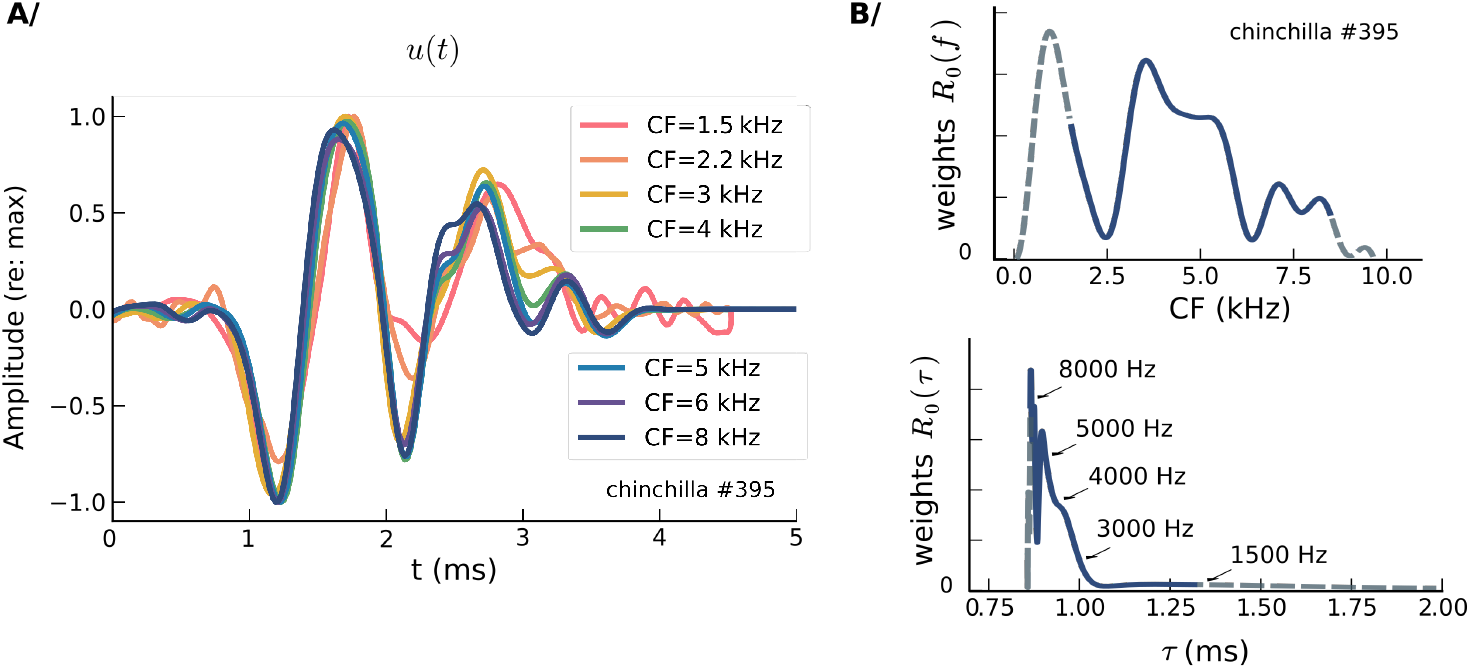
Other ancillary parameters of the model. **A**/ Estimated unitary responses *u* at the different CFs, corresponding to the weighted average of deconvolutions of Δ*CAP* responses (notched-noise maskers with varying notch attenuation) with masking release patterns. The unitary responses have been normalized according to their maximum baseline-to-peak amplitude. **B**/ Estimation of the frequency weight distribution *R*_0_(*f*) (top) representing the relative contributions of different frequencies (i.e., cochlear places) to Δ*CAP*. The weights below 1.5 kHz and above 8kHz (dashed lines) are a result of extrapolation and do not correspond to real data points. The associated distribution in the latency domain is shown (bottom). The conversion from frequency to time was done using the relation 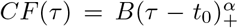, with the change of variable *R*_0_ (*f*)*df* = *R*_0_ (*f*)*B_α_*(*τ* — *t*_0_)^*α*–1^ *d_τ_* = *R*_0_ (*τ*)*d_τ_*.

### Fitting of Δ*CAP* and estimation of frequency selectivity

Figure 7 A shows how the model fit real data for two masking conditions after optimization of the model parameters (CF=1.5 kHz). The panel B in the same figure shows a synthesis of prediction errors for the same animal at all the CFs, considering all the notched-noise maskers with a notch around CF. In most cases, more than 90% of the variance was accounted for by the model. Remarkably, for some CFs, the prediction error almost reached noise level (after pre-processing). Similar accuracy numbers were obtained for the other animals in the study.

**Figure 7:**
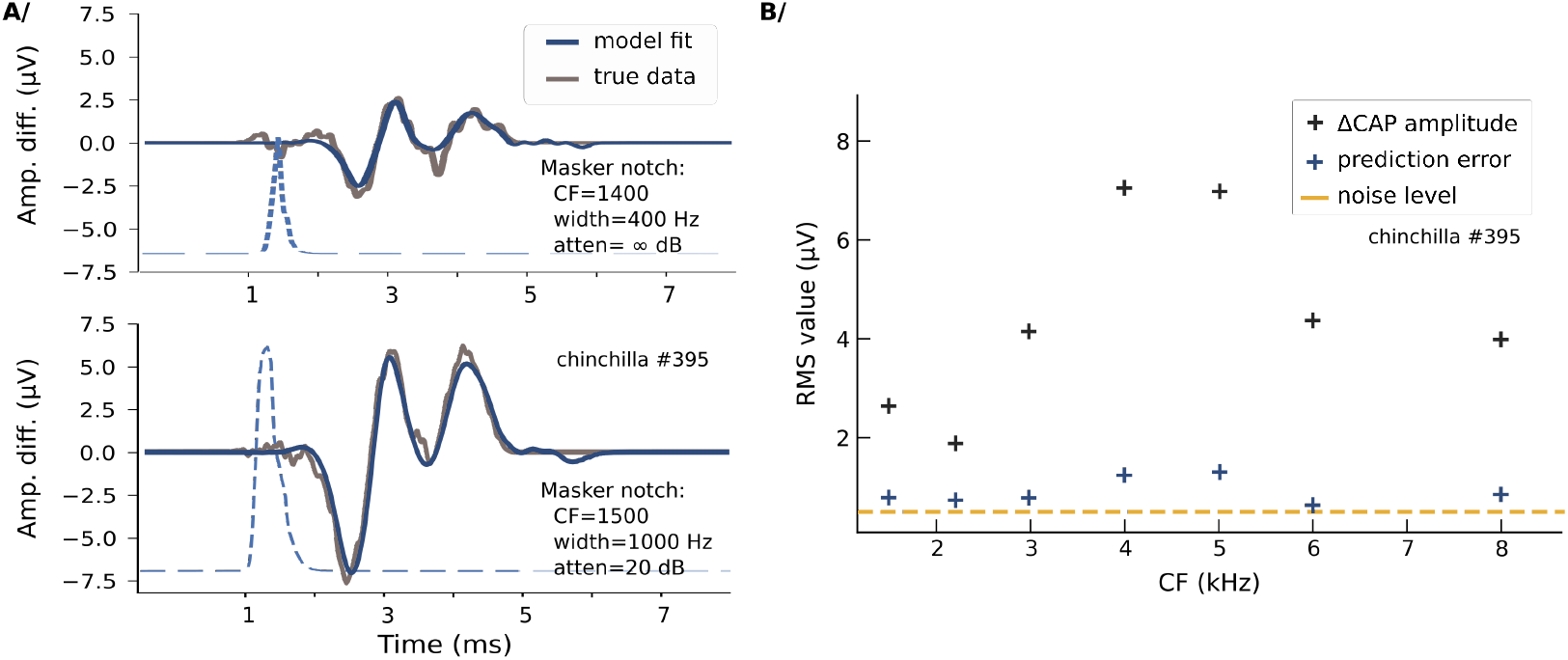
Fitting of Δ*CAP*(*t*) for one chinchilla. **A**/ Two examples of fits of Δ*CAP*(*t*) for two notched-noise maskers after parameter optimization (the first masker belongs to the varying notch width type, the second masker belongs to the varying notch attenuation type; CFs are around 1.5 kHz). Masking release excitation patterns are shown in dashed blue (arbitrary scale and zero for y-axis). **B**/ Synthesis of errors and Δ*CAP* RMS amplitude value (computed on the 100% region of the Tukey window after pre-processing of the data) at the different CFs for the same animal. The squared errors are averaged across all conditions corresponding to notched-noise maskers with a notch centered around CF.

Finally, we present the results of the estimation of frequency selectivity which was the main object of this study. Fig 8 A shows the RMS fitting error for Δ*CAP* corresponding to the maskers presenting a varying notch width around CF=5 kHz as a function of model filter bandwidth. The bandwidth minimizing the prediction error provides an estimate of the 10-dB bandwidth at CF (grid search). Alternatively, the quality factor *Q*_10_ can be optimized by gradient descent during the optimization procedure along with the other parameters of the model. We also estimated *Q*_10_ as a function of CF using a RBF network, to take advantage of the assumed regularity of the quality factor as a function of frequency. The results for all CFs and animals are presented in Fig 8 B, as well as their averages and standard deviations. An average of *Q*_10_ values derived directly from AN tuning curves is also provided for comparison, highlighting the close match between the two datasets.

**Figure 8:**
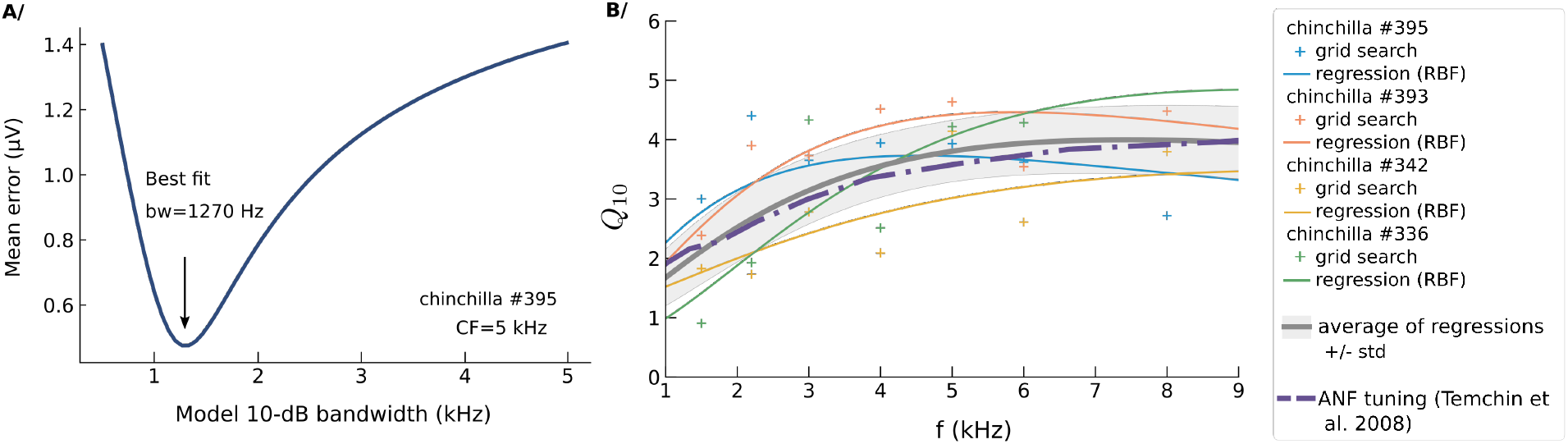
Estimation of frequency selectivity. **A**/ Grid search method: an estimate of the 10-dB bandwidth is obtained by minimizing the RMS fitting error of 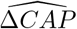 as a function of model bandwidth. In this example, the error is minimized over the responses corresponding to the notched noise maskers presenting a varying notch width around CF=5 kHz. Other parameters were considered fixed and their values were set by gradient descent before the grid search. **B**/ Synthesis of the estimates of the quality factor *Q*_10_ as a function of frequency. Crosses correspond to the estimates using the grid search method; solid lines correspond to estimates using a regression technique during the optimization procedure (RBF network). The gray shaded area shows the average and standard deviation of these solid lines. Average data from auditory nerve experiments in chinchillas [Temchin et al., 2008] are given for comparison (dashed purple line).

## Discussion

### Suitability of the convolution model for forward-masked CAPs

Our approach to fit forward-masked CAP responses with a differentiable convolution-based model led to accurate predictions of the CAP waveforms in the presence of notched Gaussian noise maskers, with more than 90% of the variance explained on the release-of-masking Δ*CAP*(*t*) in most animals and CFs. The generation of the waveform estimates relies on a consistent set of parameters, which are estimated by gradient descent (parametric I/O masking function, frequency weights, quality factor *Q*_10_) or by a specific procedure (latencies and unitary responses). A particular observation of the good performance of the model is that it was able to predict the overall shape of the release-of-masking Δ*CAP*, suggesting that the assumption that the effect of masking can be captured by a simple convolution model is valid. For instance, it is clearly apparent in Fig 3 that, if we discard the variation in amplitude, the shape of Δ*CAP* remains almost the same. We observed an exception during a pilot experiment, in which Δ*CAP* was delayed when the masking release was small (delay of 0.1 ms when the masker notch attenuation is reduced from 15 dB to 6 dB). A possible explanation for this delay is that the onset of the compound PSTH tends to be masked before the offset of the PSTH when the masker intensity is progressively increased. The probe sound level was lower for this pilot experiment than for the following sessions, suggesting that a sharper PSTH onset obtained when a higher probe level is used resolves this issue.

The latencies related to place or CF by a power-law were small above 3/4 kHz (<0.1 ms, Fig 5 B), enough to question the relevance of the convolution model for high CFs. Since all the contributions of high CFs to the masking release pattern are essentially synchronous, they could just as easily be described as a single excitation. However, the convolution model includes this particular case and can provide a more accurate model for lower CFs for which latencies are more significant. Although it captured the overall trend well, the local dependence of latencies on CF was also not always properly described by a single power-law fitted over the entire range of CFs.

The fitting of the *ΔCAP* waveforms after some adjustments of the model was remarkably accurate, but the estimation procedure presented several challenges. The fact that the model relies on a relatively large number of parameters, especially if we include all the possible dependencies on CF, can make the optimization cumbersome. The optimization of the model is however facilitated by the existence of new elegant libraries for automatic differentiation. We found that the main difficulty regarding the estimation of the different model parameters was the determination of the weights *R*_0_(*f*). The model would be greatly simplified if we could assume that the contributions to Δ*CAP* are homogeneous across CFs, but we found that it was not the case. We showed one extreme case in Fig 6 B where two narrow dips (at 2.5 kHz and 6 kHz) are present. The estimation of *R*_0_(*f*) is still possible with regularity assumptions and notched-noise maskers with notches distributed over the entire range of frequencies. However, if the dips are too steep, the estimation of the frequency weights and of the other parameters can be affected. As potential evidence, the largest deviation between *Q*_10_ values derived from AN tuning curves and those obtained with our estimation procedure (Fig 8 B) was observed at 2.2 kHz for the animal presenting a dip around this frequency (grid search method). By using a regression technique for the estimation of *Q*_10_, we can however exploit the regularity of the quality factor with respect to CF to still provide an accurate estimate of frequency selectivity (solid lines in Fig 8 B).

### Estimation of frequency selectivity using forward-masked CAPs

We found a good agreement between the estimates of the quality factor averaged over the 4 experiments for which we had complete data (Fig 8) and published values derived from the collection of many ANF tuning curves in chinchillas[Temchin et al., 2008]. Our experimental approach that led to the estimation of cochlear frequency selectivity was inspired by the experiments of Verschooten et al. [Verschooten et al., 2012, 2018] – their work was in turn an improvement of experiments using forward-masked CAPs that were conducted in the 1980s [Harrison et al., 1981a,b]. The method of frequency selectivity estimation used by Verschooten et al involved establishing iso-response curves for masker level versus masker notch width – the response criterion being that 66% of the initial CAP amplitude had to be restored. A measure of tuning was derived from these curves by considering the 10-dB bandwidth – reduced to a single auditory filter model, this measure can be seen as the bandwidth encompassing 90% of the frequency response power spectrum (called BW90 in other works [Unoki et al., 2006]). The main interest of their technique compared to ours is that it does not require the assumption that the amount of masking of synchronized ANFs is driven by input-output curves that are to be determined. Rather, their measure of tuning was considered as an empirical quantity, and assumed to be proportional to the 10-dB bandwidth of ANF tuning curves. They found a good agreement between the two quantities after a constant correction factor was applied. However, the conversion factor from CAP to ANF data was not the same for every species and smaller for small mammals. In addition, the correction factor for macaques was not constant as a function of frequency (S5 Fig in [Verschooten et al., 2018]). It is therefore not clear how the derived measure can be interpreted, as it may be affected differently from one species to another by, for example, the effect of off-frequency masking. The method presented in this article provides a more direct estimate of frequency selectivity that is grounded by a mathematical model of forward-masked CAPs and does not require an empirical correction factor. The convolution-based method has other advantages. Since the entire *ΔCAP* signal is used instead of the CAP peaks, the estimation is more robust to noise. Furthermore, it exploits all the available data, whereas the ‘fast’ procedure in Verschooten et al. searches for a particular masker level meeting the masking criterion, thus potentially wasting measurements points. Beyond these aspects, a potential of our method is that the mathematical model and experimental approach could be adapted to study more complex aspects of cochlear signal processing, such as compressive nonlinearities, as explained in the next paragraph.

### Limitations related to the simplified underlying auditory model

A few difficulties associated with the model were mentioned throughout the paper, including changes in the model parameters with CF that make the estimation more challenging. Another set of limitations is related to the oversimplifications of the model to describe the behavior of the cochlea. The main shortcoming of the model is that the cochlear decomposition of the signal, which was merely described as the action of a filter bank (Fig 2), is in reality not linear. Compressive nonlinearities not only affect frequency tuning by broadening the auditory filters when intensity is increased [Heinz et al., 2002], but they also modify the input-output functions depending on the amount of suppression [Delgutte, 1990]. Therefore, including these nonlinear effects in the model represents a major challenge for future developments, but could lead to a detailed picture of how compressive nonlinearities affect cochlear processing. To study nonlinear effects properly, a greater variety of masking conditions would also have to be employed during data collection. As an example, Verschooten et al. evaluated the level dependence of cochlear frequency selectivity in cats by presenting maskers of various intensities [Verschooten et al., 2012].

Other aspects of the auditory model considered in this work correspond to oversimplifications of cochlear signal processing. Auditory filter frequency profiles are in reality asymmetric, and the lower and upper sides are not affected the same way by nonlinearities [Irino and Patterson, 2002]. We focused on the ‘tip’ of the auditory filters, which can be accurately described by gammatones or Gaussian filters – and we did not find significant differences using one model or the other – but auditory filters also present a low-frequency tail, the latter showing different attributes depending on filter CF [Temchin et al., 2008]. In addition, the preferred frequency of auditory filters change with the degree of compression [Lopez-Poveda and Meddis, 2001]. Future work is needed to explore whether the proposed method could be extended to include these nonlinear properties and prove to be an effective tool to study complex aspects of cochlear signal processing with a lower degree of invasiveness compared to single-fiber AN recordings.

## Supporting information

SI 2: graph of computations

SI 1: Compound PSTH (Simulation)

## Acknowledgments

This work was conducted while the first author was on a postdoctoral fellowship supported by Fondation Pour l’Audition (FPA RD-2019-3). Support was also provided by NIH grants R01-DC009838 and T32-DC016853.

## Appendix

### Computation of cochlear-filter output intensity for the gammatone model

*Note: In this paragraph, τ does not have the same use as in the main part of the paper where it is a variable for latencies. Here, it refers to the time constant of the gammatones.*

The k-th order gammatone, characterized by an envelope proportional to 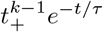, is defined in the frequency domain (complex version, w.l.o.g.) by:

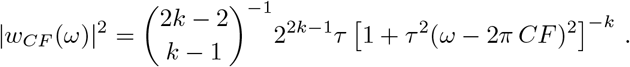

Considering one band of noise masker (the results simply add up if there are multiple bands), we have:

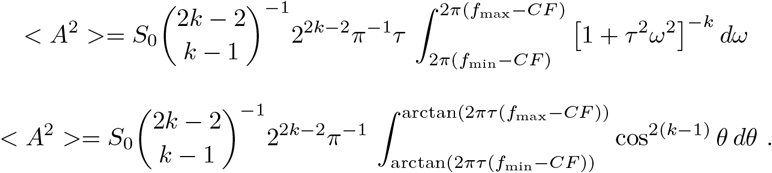

The last integral is then computed by writing

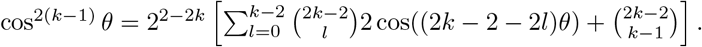

*Note: The 10-dB bandwidth is related to *τ* by* 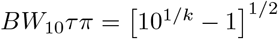.

## Notes

https://zenodo.org/record/6403024

### Competing Interest Statement

The authors have declared no competing interest.

