## Supplementary material for "Estimation of cochlear frequency selectivity using a convolution model of forward-masked compound action potentials": SI 2: graph of computations

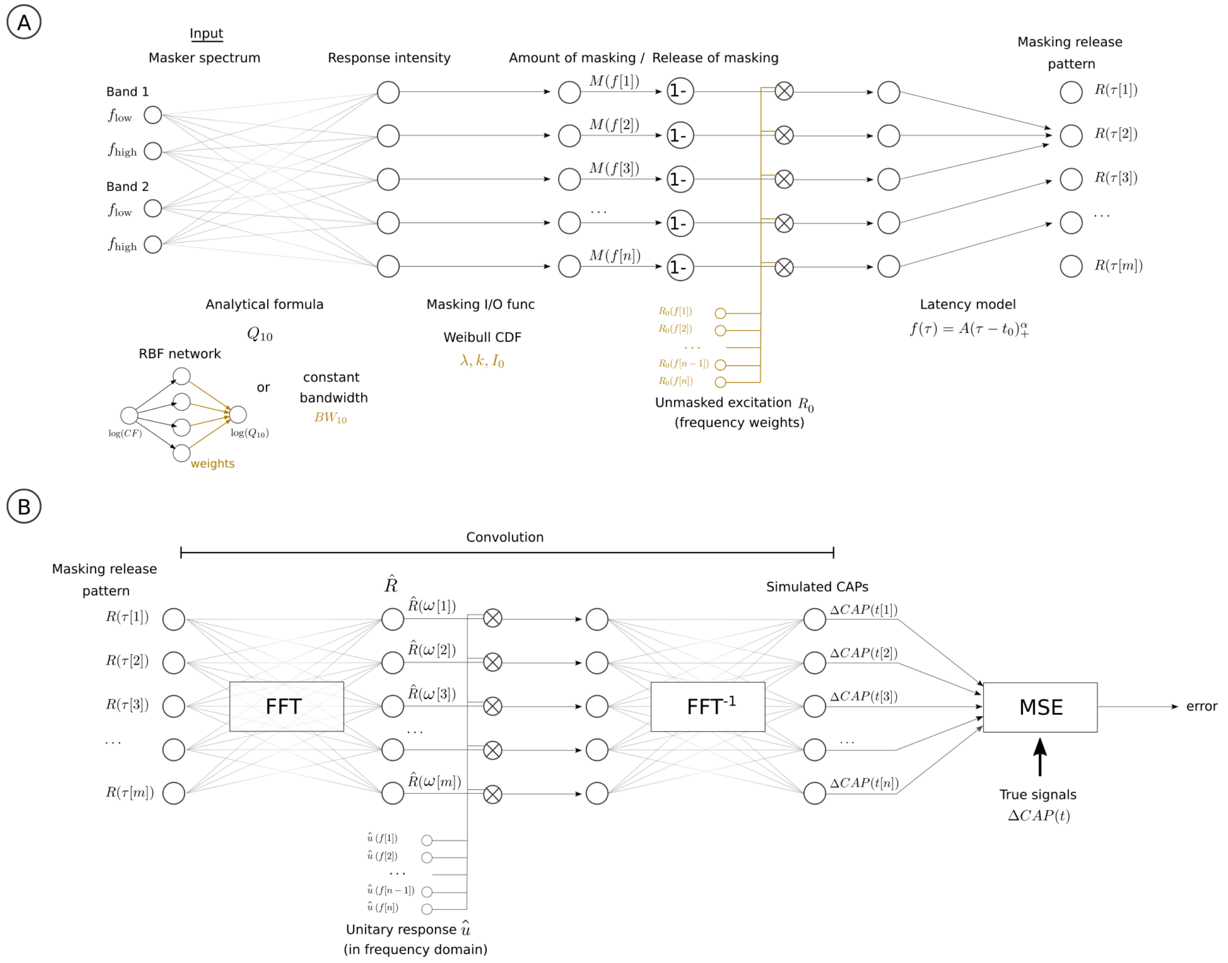


Figure 1: Diagram of computations (divided in two parts A and B) for a masker with 2 bands implemented using PyTorch leading to the generation of the estimated CAP release-of-masking $\widehat{\Delta CAP}(t)$.The variables that are updated during the optimization procedure (gradient descent) are represented in gold font. Analytical formula for response intensity: see main text for its mathematical expression.
