## Supplementary material for "Estimation of cochlear frequency selectivity using a convolution model of forward-masked compound action potentials": SI 1: Compound PSTH (Simulation)

**Simulation of masking on the compound response of auditory nerve fibers (ANFs)**

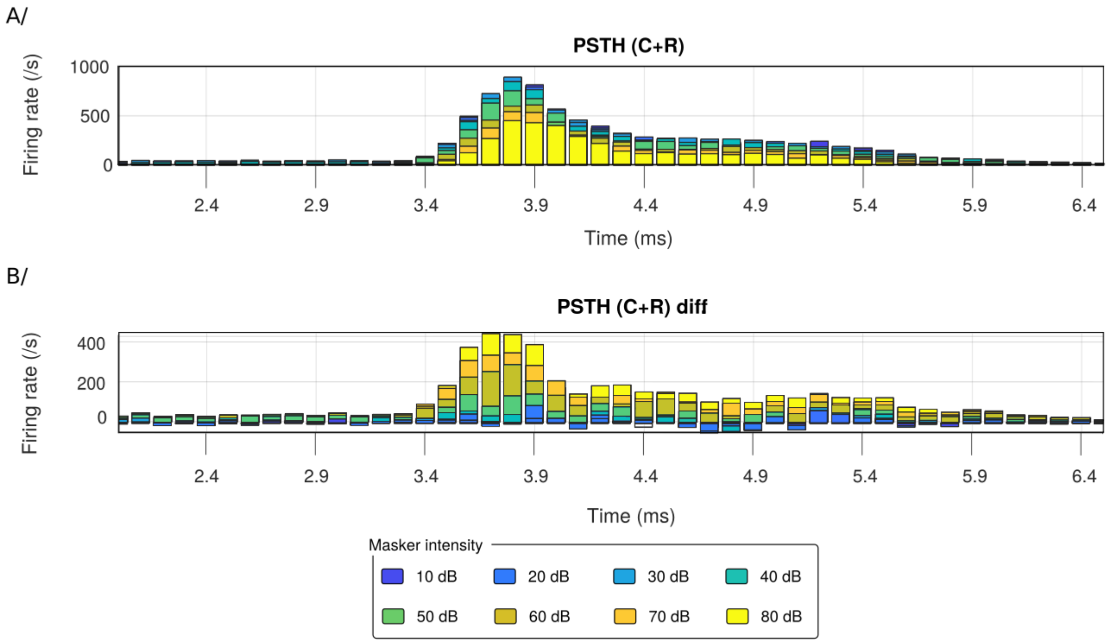


**A/** Post-stimulus time histograms (PSTHs) of a population of ANFs using the Bruce et al. model (2018). The bar plots are the compound PSTHs in response to a 90 dB SPL click presented 5 ms after a 30-ms white Gaussian noise masker with a masker level varying from 10 dB to 80 dB. Simulation parameters: CF = 4 kHz, population of 32 ANFs (low spontaneous rate LS: 6, MS: 6, HS: 20); N reps = 400 (clicks were of alternating polarity); bin interval: 0.1 ms. **B/** Difference in the compound PSTHs taking as reference the response to masker with the lowest level. This figure shows that the PSTH differences present a similar shape when normalized with respect to the amount of masking or the total number of spikes. This observation was in favor of the use of the convolution model for $\Delta CAP(t)$. When normalized, the PSTH differences play the role of $n\Delta PST$ in the main text.
